# The hourglass model of evolutionary conservation during embryogenesis extends to developmental enhancers with signatures of positive selection

**DOI:** 10.1101/2020.11.02.364505

**Authors:** Jialin Liu, Rebecca R. Viales, Pierre Khoueiry, James P. Reddington, Charles Girardot, Eileen E. M. Furlong, Marc Robinson-Rechavi

## Abstract

Inter-species comparisons of both morphology and gene expression within a phylum have revealed a period in the middle of embryogenesis with more similarity between species compared to earlier and later time-points. This ‘developmental hourglass’ pattern has been observed in many phyla, yet the evolutionary constraints on gene expression, and underlying mechanisms of how this is regulated, remains elusive. Moreover, the role of positive selection on gene regulation in the more diverged earlier and later stages of embryogenesis remains unknown. Here, using DNase-seq to identify regulatory regions in two distant *Drosophila* species (*D. melanogaster* and *D. virilis*), we assessed the evolutionary conservation and adaptive evolution of enhancers throughout multiple stages of embryogenesis. This revealed a higher proportion of conserved enhancers at the phylotypic period, providing a regulatory basis for the hourglass expression pattern. Using an *in silico* mutagenesis approach, we detect signatures of positive selection on developmental enhancers at early and late stages of embryogenesis, with a depletion at the phylotypic period, suggesting positive selection as one evolutionary mechanism underlying the hourglass pattern of animal evolution.

## Introduction

Embryological development has long been characterized by deep conservation, from morphology to mechanisms. Animals that belong to the same phylum share a group of structural and developmental characteristics, the so called basic body plan or bauplan (Wallace 2000; Valentine 2004; Irie and Kuratani 2014). For example, arthropods share a set of anatomic structures such as jointed legs, an exoskeleton made of chitin, and segmented bodies (Zrzavý and Štys 1997). Based on morphological conservation, Duboule (1994) and Raff (1996) proposed the hourglass model. Under this model, within a phylum, embryos at mid-embryonic stages (the phylotypic period (Richardson 1995)) are morphologically more conserved than embryos in both early and late development. However, this model was not supported by later morphological studies (Richardson et al. 1997; Bininda-Emonds et al. 2003). To overcome difficulties of comparing morphological features across species, more recent studies used comparative transcriptomics (Yanai 2018), as changes in gene expression play a central role in the morphological differences between species (King and Wilson 1975; Carroll 2008).

Transcriptome comparisons in different phyla (Kalinka et al. 2010; Irie and Kuratani 2011; Levin et al. 2012; Schep and Adryan 2013; Hu et al. 2017) indicate that expression divergence is lower in the phylotypic period compared to early and late stages of embryogenesis, supporting the hourglass model. One of the pioneer studies was conducted in six *Drosophila* species by Kalinka et al. (2010), quantifying expression divergence at different stages during embryogenesis. They found that expression divergence follows an hourglass pattern with the minimum divergence at the extended germband stage (8-10 hours after laying egg), generally regarded as part of the arthropod phylotypic period (Sander 1976). Notably, this expression hourglass pattern also extends to the population level (Zalts and Yanai 2017), and even to the level of variation between isogenic individuals (Liu et al. 2020). Based on a single embryo transcriptome time series of *Drosophila* embryonic development, with a high number of isogenic replicates, we found that the phylotypic period also has lower non-genetic expression variability (Liu et al. 2020).

Despite many transcriptomic comparisons, the role of the underlying regulatory regions (e.g. developmental enhancers) on the evolution of expression during embryogenesis remains to be elucidated. What’s more, although purifying selection and mutational robustness can explain the hourglass expression divergence pattern (Zalts and Yanai 2017; Liu et al. 2020), the contribution of positive selection to the hourglass model remains unknown. For example, this pattern may also result from enhanced positive selection at both the early and late development stages. Moreover, the two evolutionary mechanisms (purifying versus positive selection) may not be mutually exclusive. In terms of protein sequence, for example, the lower sequence evolution in the phylotypic period appears to be caused by both strong purifying selection and weak positive selection (Liu and Robinson-Rechavi 2018a; Liu and Robinson-Rechavi 2018b; Coronado-Zamora et al. 2019). To investigate the underlying regulatory mechanisms, and the contribution of positive selection at regulatory elements, to the hourglass model, we performed DNase-seq to identify active regulatory elements across multiple matched embryonic developmental stages in two distant *Drosophila* species: *D. melanogaster* and *D. virilis*.

## Results

### DNase-seq across five stages of embryogenesis in two species

To study the evolution of enhancers in the context of embryonic development, we extended our previously published DNase I hypersensitive sites sequencing (DNase-seq) data on three embryonic stages of *D. virilis* and *D. melanogaster* (Peng et al. 2019) to five equivalent embryonic stages in both species (TP1 to TP5, Fig. 1A). TP3 is part of the phylotypic period (Fig. 1A), with the two new time points extending to later stages beyond the phylotypic period. Regulatory regions were identified using DNase I hypersensitive sites sequencing (DNase-seq) in tightly staged whole embryos. Every stage had two biological replicates from each species, and high confidence peaks (bound regulatory regions) were called at a 5% Irreproducible Discovery Rate (IDR, a measure ensuring equivalent reproducibility between replicas (Li et al. 2011)). On average, we identified 15,831 peaks in each stage in *D. virilis*, and 14,995 peaks in *D. melanogaster* ((Supplemental Fig. S1). Replicates are highly concordant both for raw reads (median Spearman’s correlation coefficient 0.96 for *D. virilis*, 0.92 for *D. melanogaster*) and for significant peaks (median Spearman’s correlation coefficient = 0.94 for *D. virilis*, 0.90 for *D. melanogaster*). Distal regulatory elements, including putative enhancers, were defined as peaks greater than 500bp from an annotated transcriptional state site (TSS). While these regions may also include other regulatory elements such as insulators, for the sake of simplicity, we refer to these putative enhancers simply as enhancers in the rest of the study. We detected more distal elements (enhancers) in *D. virilis* than *D. melanogaster* (Fig. 1B), similar to our previous findings using ChIP-seq against transcription factors (Khoueiry et al. 2017), which likely reflects the larger size of the *D. virilis* non-coding genome. In addition, in both species, we found that TP3 has more enhancers than other stages (Fig. 1B). The general trend in the relative number of enhancers across development is consistent between the two species.

**Figure 1:**
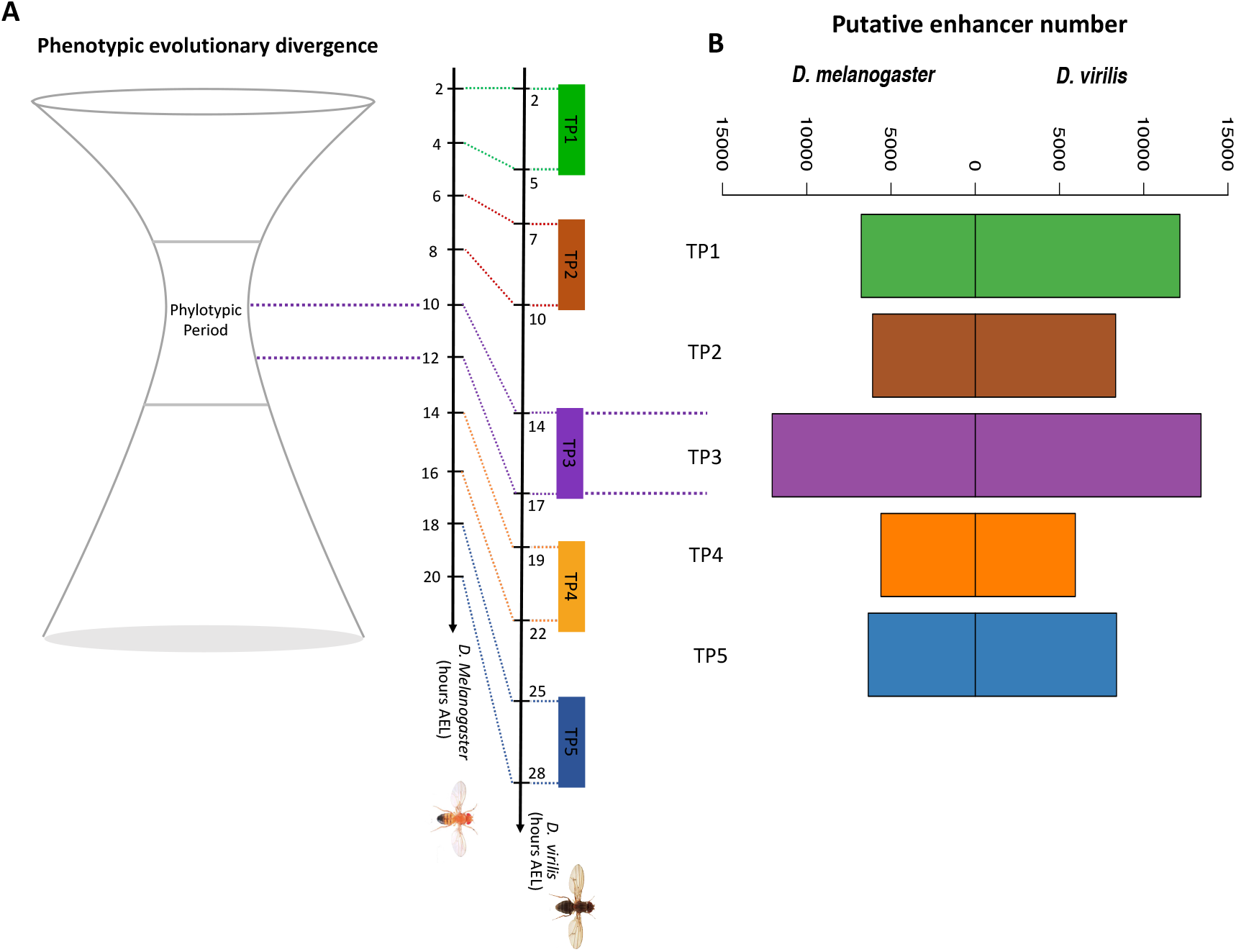
Studying regulatory elements evolution throughout embryogenesis. A. *We performed DNase-seq at two matched late embryonic development stages in D. melanogaster and D. virilis (TP4 and TP5) and combined this with our previous DNase I hypersensitive sites (DHS) data (TP1-3) in both species. The corresponding time points (TP1, TP2, TP3, TP4 and TP5) for the five embryonic stages are shown on the developmental axis. Different color bars represent different time points sampled. The developmental axes are scaled in hours after egg laying (AEL).* B. *Number of putative enhancers in the five embryonic development stages in each species.*

### Enhancer conservation over embryonic development

To compare evolutionary conservation of enhancers between stages, we first identified stage-specific enhancers for each time-point in each species. For example, *D. melanogaster* TP3 stage-specific enhancers were defined as regions with a DNase peak in this stage and no significant peak at other stages in *D. melanogaster* (Fig. 2A; Table 1). For all stage-specific enhancers in one species, we identified their corresponding orthologous regions in the other species with pslMap (Zhu et al., 2007) (see Methods), restricting to one-to-one orthologous regions. In both species, we found that TP3, within the phylotypic period, has a higher proportion of enhancers with orthologous regions (Supplemental Fig. S2), indicating stronger sequence conservation for phylotypic period-specific enhancers. Next, we identified conserved stage-specific enhancers. For example, if a *D. melanogaster* TP3 specific enhancer has both an orthologous region and an overlap (by orthologous translation) with a *D. virilis* TP3 specific enhancer, we defined this as a conserved TP3 specific enhancer (Fig. 2A). Finally, to quantify the overall conservation, at each stage, we calculated the Jaccard index (Fig. 2B), which ranges from 0 (none of the stage-specific enhancers are conserved) to 1 (all stage-specific enhancers are conserved). We found that TP3 has a significantly higher proportion of conserved enhancers than other stages (Fig. 2C). Given the larger size of non-coding genome in *D. virilis*, the genome coordinates of orthologous enhancers can be shifted by insertions and deletions. To account for this, we repeated the analysis with a relaxed definition of conserved enhancer: the distance between stage-specific enhancers in the two species was defined to be smaller than 1kb, but not necessarily overlap. We found a very similar pattern with this more relaxed definition, with a higher proportion of conserved enhancers at TP3 (Fig. 2D). In addition, similar patterns were observed when we used all enhancers, not restricting to stage specific enhancers (Supplemental Fig. S3).

**Figure 2:**
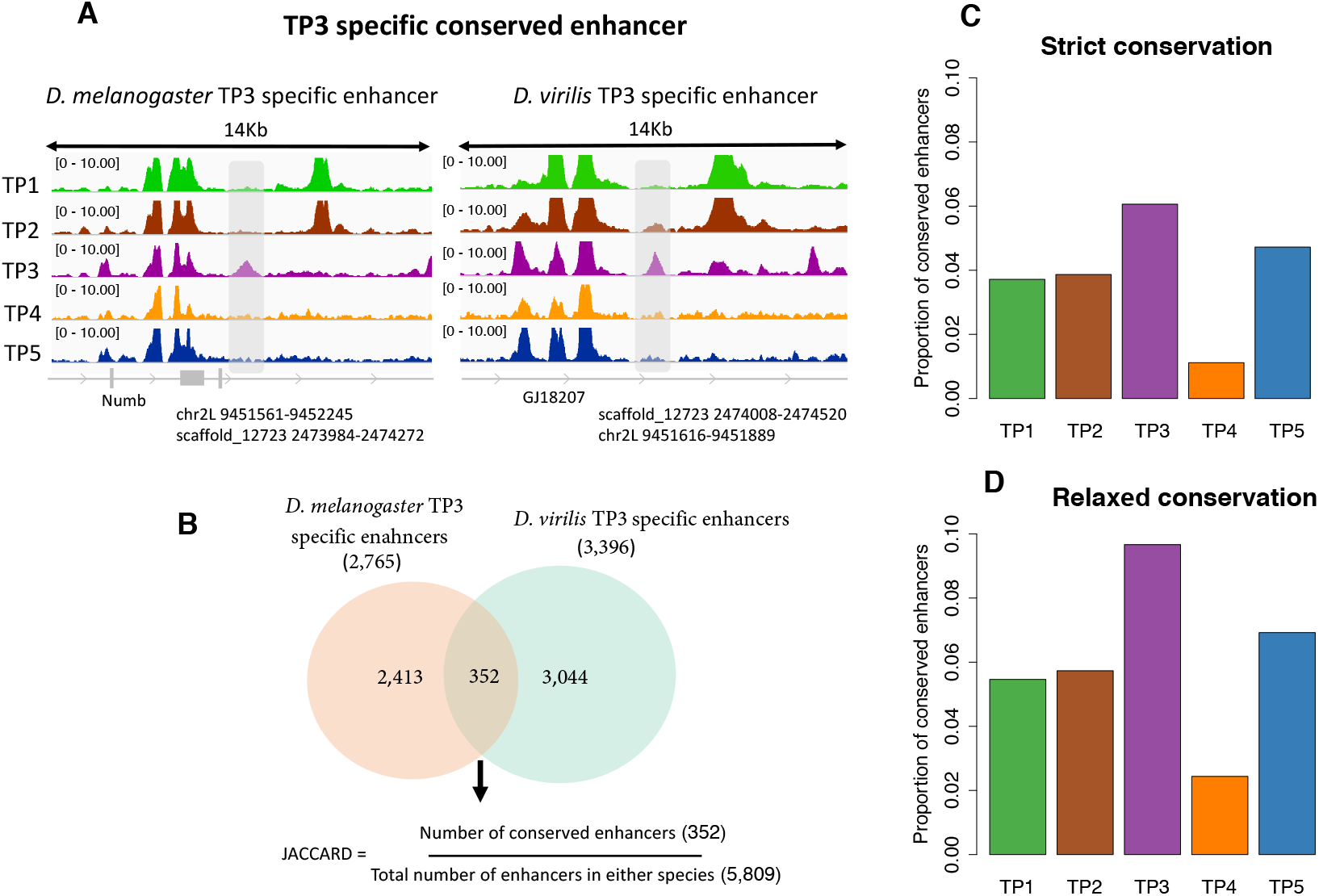
The phylotypic period has a higher proportion of conserved regulatory elements. A. *Illustration of a TP3 specific conserved enhancer. The left panel is the DNase-seq signal in different development stages (TP1, TP2, TP3, TP4 and TP5) for a D. melanogaster TP3 specific enhancer, covered by the grey area. The coordinates of this enhancer in D. melanogaster, and the orthologous coordinates in D. virilis, are indicated below the grey arrow. The right panel is the DNase-seq signals in different development stages (TP1, TP2, TP3, TP4 and TP5) for a D. virilis TP3 specific enhancer, covered by the grey area. The coordinates of this enhancer in D. virilis, and the orthologous coordinates in D. melanogaster, are illustrated below the grey arrow. Since there is overlap between the D. melanogaster TP3 specific enhancer and the D. virilis TP3 specific enhancer based on orthologous position, we define the two enhancers as a TP3 conserved enhancer.* B. *Venn diagram of orthologous TP3 specific enhancers (only one-to-one orthologs) conserved between both species.* C. *Proportion of conserved stage-specific enhancers at each development stage. Here the conservation means there is at least 1bp overlap between stage-specific enhancers in the two species. The p-values from pairwise Fisher’s exact tests between TP3 and TP1, TP2, TP4, and TP5 are 6.51 × 10^−7^, 3.69 × 10^−3^, 1.44 × 10^−14^, 2.22 × 10^−2^ respectively.* D. *Proportion of conserved stage-specific enhancers at each development stage. Here the conservation means the distance between stage-specific enhancers in the two species must be smaller than 1kb, not necessarily overlap. The p-values from pairwise Fisher’s exact tests between TP3 and TP1, TP2, TP4, TP5 are 1.89 × 10^−12^, 4.28 × 10^−5^, 2.7 × 10^−17^, 2.25 × 10^−4^ respectively.*

**Table 1:**
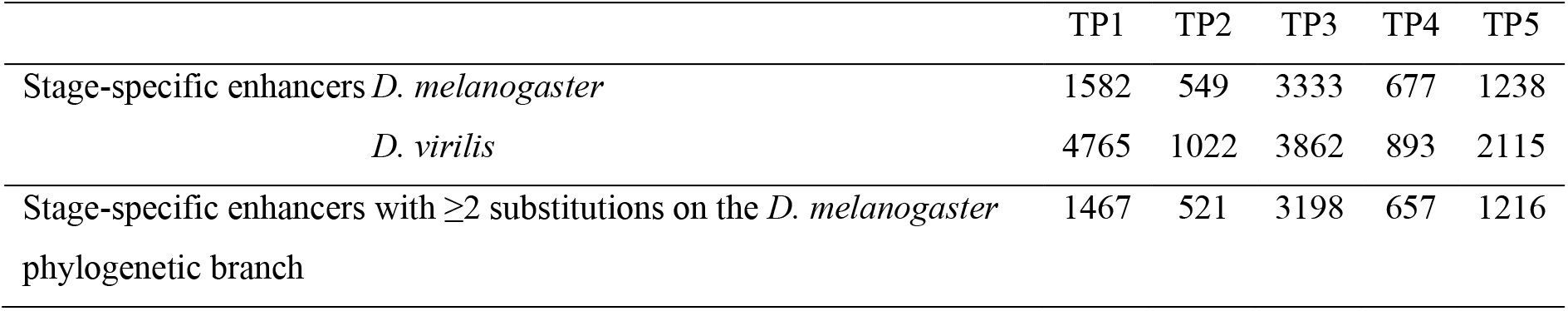
Number of enhancers for each sampled stage.

|  | TP1 | TP2 | TP3 | TP4 | TP5 |
| --- | --- | --- | --- | --- | --- |
| Stage-specific enhancers *D. melanogaster* | 1582 | 549 | 3333 | 677 | 1238 |
| *D. virilis* | 4765 | 1022 | 3862 | 893 | 2115 |
| Stage-specific enhancers with $\geq 2$ substitutions on the *D. melanogaster* phylogenetic branch | 1467 | 521 | 3198 | 657 | 1216 |

### Detecting positive selection on enhancers

Higher conservation can be explained either by stronger purifying selection or by weaker positive selection. To test for the latter, we scanned for signatures of positive selection in all *D. melanogaster* stage-specific enhancers. Our approach considers the effects of substitutions on enhancer accessibility, and is derived from a new method to detect positive selection in transcription factor binding site evolution (Liu and Robinson-Rechavi 2020). Briefly, a gapped k-mer support vector machine (gkm-SVM) classifier is trained on these stage-specific enhancers. This gkm-SVM identifies sequence features that determine chromatin accessibility, thus enhancer occupancy. This allows computing SVM weights of all possible 10-mers, which are predictions of their contribution to enhancer accessibility. We can then predict the accessibility impact of substitutions by calculating deltaSVM, the difference of sum weights between two homologous sequences. The significance of the observed deltaSVM was evaluated by comparing it with a null distribution of deltaSVM, constructed by scoring the same number of random substitutions 10,000 times. The p-value can be interpreted as the probability that the observed deltaSVM could arise by chance under the assumptions of the randomization.

Adaptive evolution on enhancer chromatin accessibility is expected to push them from a suboptimal accessibility toward an optimal accessibility, or from an old optimum to a new one. Thus, chromatin accessible sites evolving adaptively are expected to accumulate substitutions that consistently change the phenotype towards stronger or towards weaker accessibility, whereas sites evolving under relaxed purifying selection are expected to accumulate substitutions that increase or diminish accessibility randomly around a constant optimum.

As this positive selection scanning method needs to be applied to sequences with relatively low divergence, we tested for substitutions on the *D. melanogaster* branch after divergence from *D. simulans* (Fig. 3A), rather than the much more distant *D. virilis* branch. To check whether there were sequence features which differ between enhancers of different stages, we first separately trained a gapped *k*-mer support vector machine (gkm-SVM) for each stage (see Methods). Not only can the gkm-SVM trained in the corresponding stage accurately distinguish enhancers from random sequences, it also has higher performance than the gkm-SVM trained from other stages. (Fig. 3B, see Methods). In addition, a gkm-SVM trained from an adjacent stage has higher performance than a gkm-SVM trained from a distant stage. For example, the gkm-SVM model trained from TP2 has higher power to distinguish TP1 specific enhancers from random sequences than the gkm-SVM model trained from TP5. These results suggest that the gkm-SVM’s predictions are not only informative, but also developmental stage specific.

**Figure 3:**
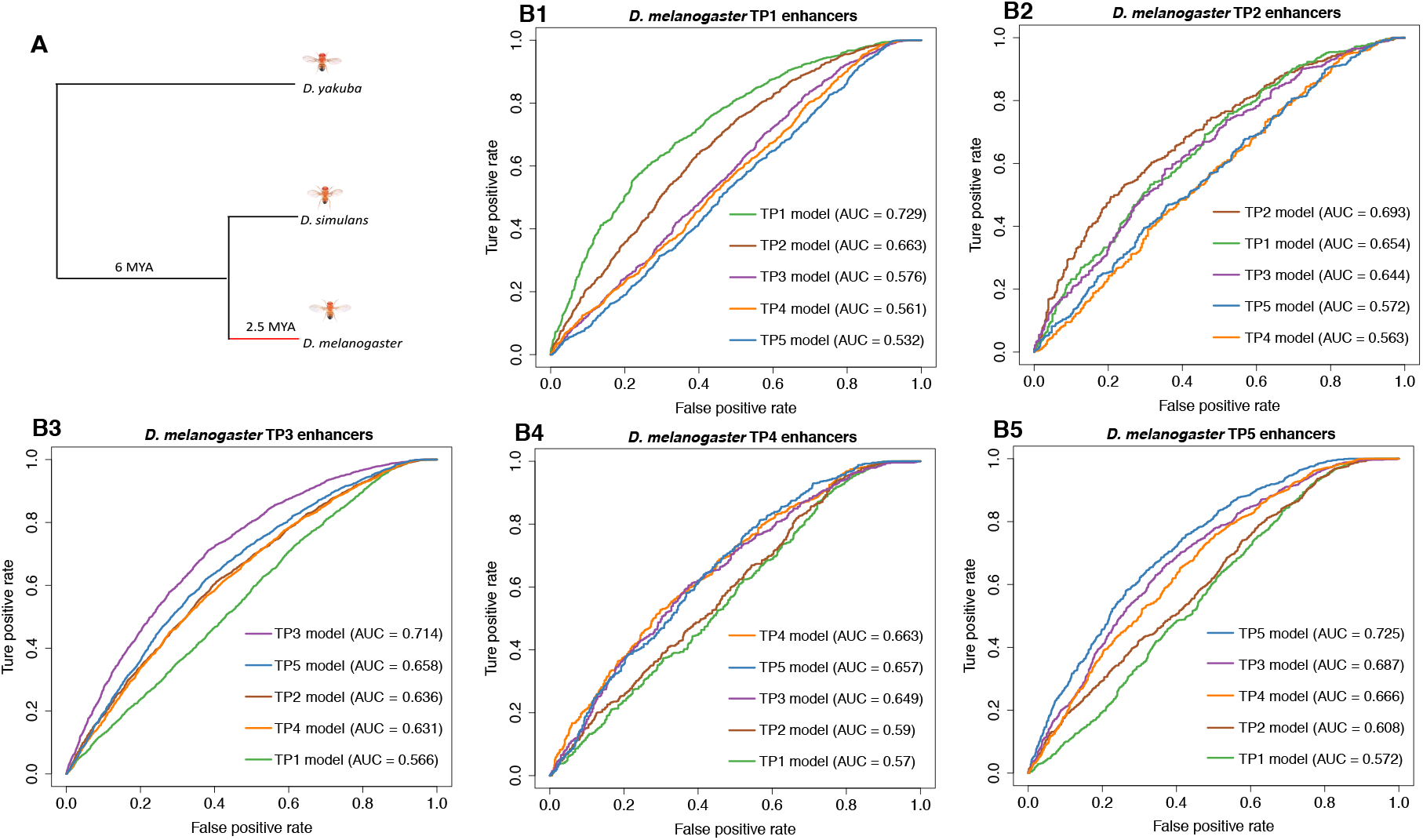
Gapped k-mer support vector machine (gkm-SVM) can predict stage-specific enhancers. A. Topological illustration of the phylogenetic relationships between the three Drosophila species used to detect positive selection on enhancers. We want to detect positive selection which occurred on the lineage of D. melanogaster after divergence from D. simulans, as indicated by the red branch. D. yakuba is the outgroup used to infer enhancer sequence in the ancestor of D. melanogaster and D. simulans. B1-5. Receiver operating characteristic (ROC) curves for gkm-SVM classification performance on stage-specific enhancers. AUC values represent areas under the ROC curve and provide an overall measure of predictive power.

We focused on enhancers with at least two substitutions to their sequence between species (Table 1). In each stage, we calculated the deltaSVM score of every stage-specific enhancer based on its corresponding gkm-SVM. We associated a *p-*value for each enhancer, by *in silico* mutagenesis (see Methods). Thus, we can identify enhancers whose substitution pattern on the *D. melanogaster* branch has effects on chromatin accessibility which are inconsistent with neutrality, and therefore imply positive selection on enhancers. In each stage, the distributions of *p-*values for all stage-specific enhancers shows a strong skew toward low *p-*values (Fig. 4A), indicating evidence for positive selection. For all downstream analyses, we use *q* < 0.05 (i.e. 5% false positives) as a threshold to define an enhancer as having evolved under positive selection (hereafter “positive selection enhancer“). Since mutations under positive selection will spread through a population rapidly, they are expected to decrease polymorphisms (intra-species variation) while increasing substitutions (inter-species variation) (McDonald and Kreitman 1991). Thus, we expect that positive selection enhancers should have higher substitutions to polymorphisms ratios than non-positive selection enhancers. To test this, we counted the number of substitutions between *D. melanogaster* and *D. simulans*, and number of polymorphisms among 205 *D. melanogaster* inbred lines from wild isolates, separately for positive selection and non-positive selection enhancers (see Methods). As predicted, in all stages, positive selection enhancers have a significant excesses of fixed nucleotide changes (Fig. 4B), confirming that we are indeed detecting positive selection in enhancer evolution.

**Figure 4:**
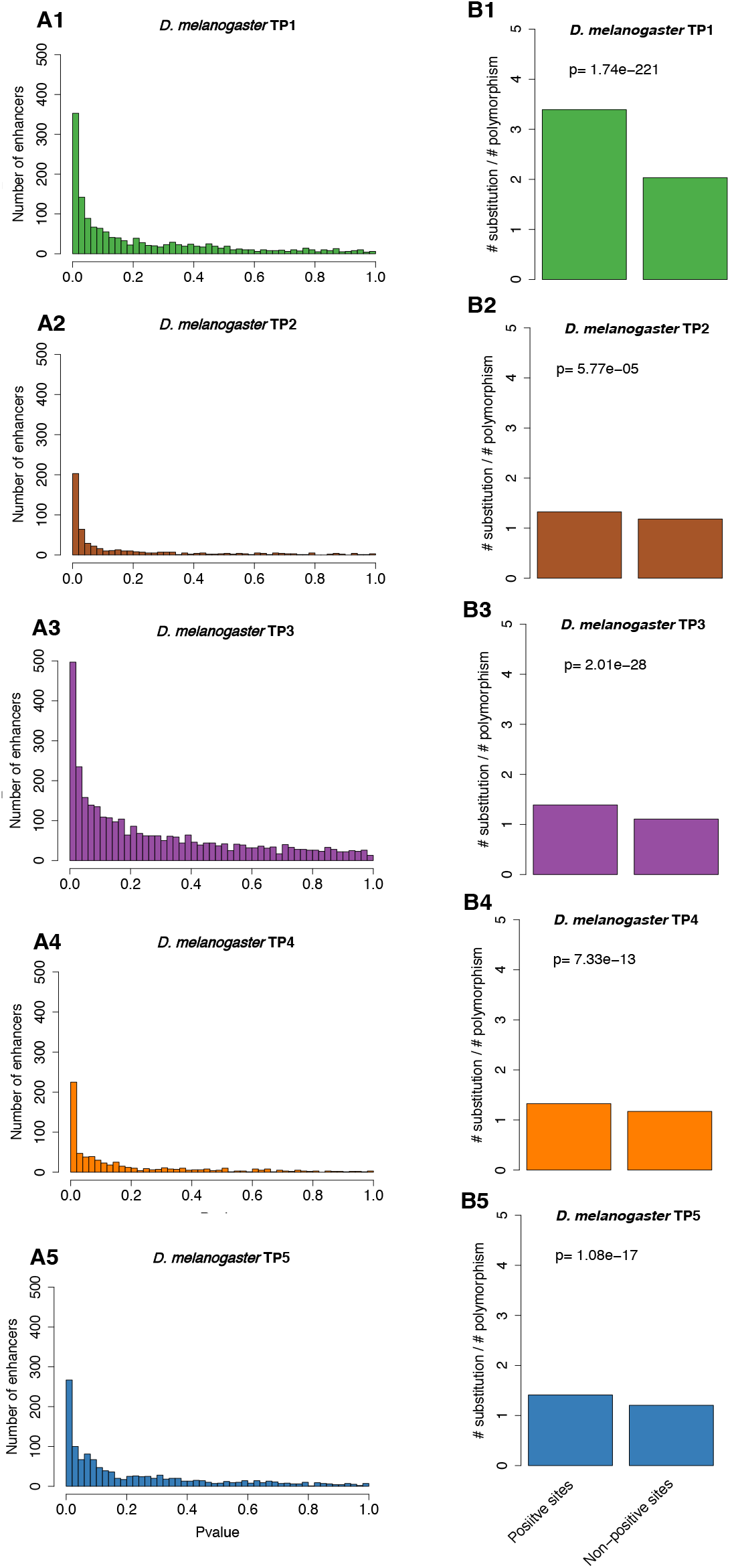
Evidence of positive selection throughout embryogenesis. A1-5. The distribution of deltaSVM p-values (test for positive selection) for each stage-specific enhancer. B1-5. The ratio between the number of substitutions and the number of polymorphisms (SNPs) for each stage-specific enhancers. Positive sites are enhancers with evidence of positive selection (deltaSVM qvalue < 0.05), non-positive sites are enhancers without evidence of positive selection. The p-value from Fisher’s exact test is reported above the bars.

Having identified enhancers which evolved under positive selection, we investigated whether their distribution varies across development. We previously found that species specific gains in transcription factor binding sites have a higher proportion of positive selection than conserved ones (Liu and Robinson-Rechavi 2020). Thus, we separated *D. melanogaster* enhancers into conserved and non-conserved enhancers, based on conservation with *D. virilis* enhancers. As expected, the non-conserved enhancers generally have a higher proportion of positive selection than the conserved ones (Fig. 5). Moreover, over development, the phylotypic period has a much lower proportion of enhancers with evidence of positive selection than the other stages (Fig. 5). This suggests that positive selection contributes to the evolution of enhancers, and that the phylotypic period is characterized by less positive selection.

**Figure 5:**
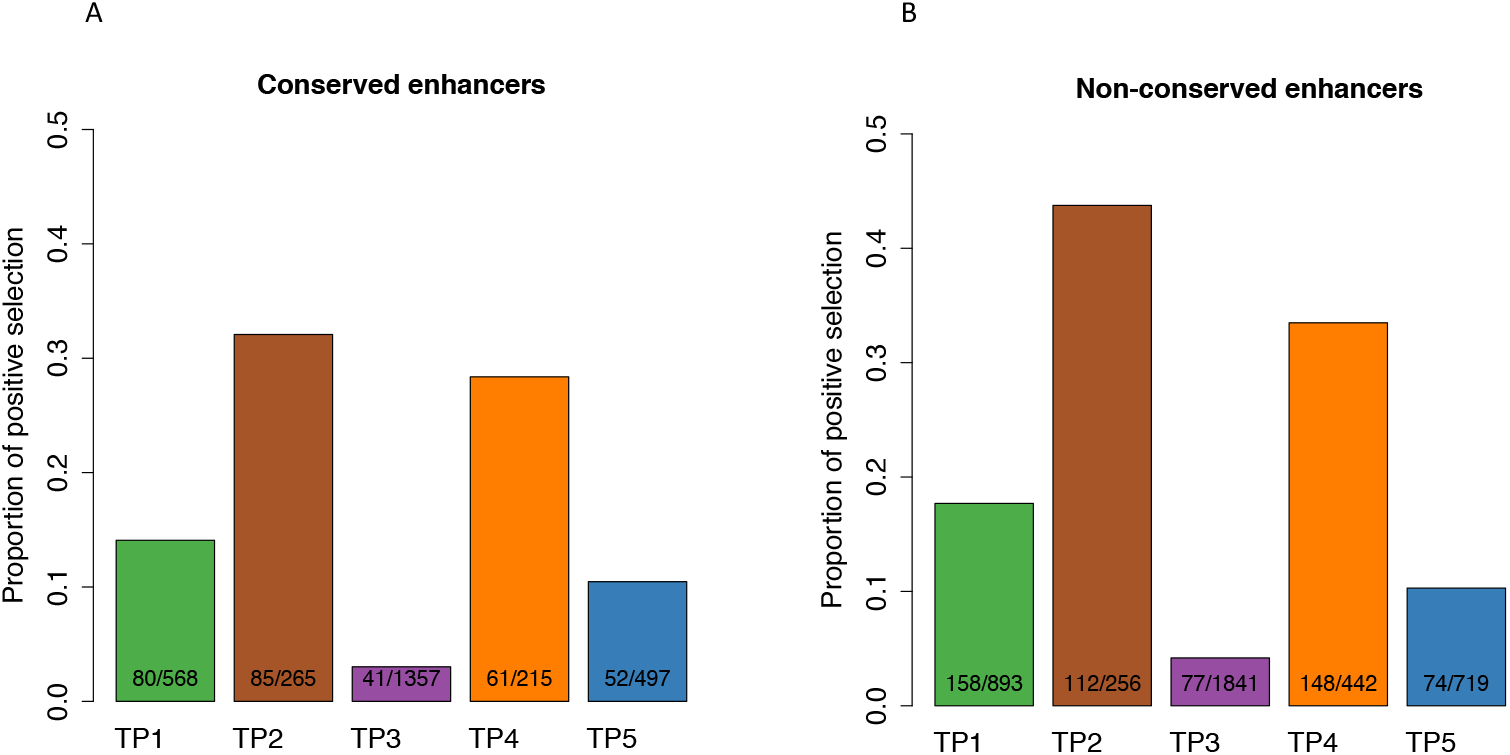
The phylotypic period has a lower proportion of enhancers with evidence of positive selection. The proportion of enhancers with evidence of positive selection in the five stages. Positive sites are enhancers with evidence of positive selection (deltaSVM qvalue < 0.05). The number of stage-specific enhancers and the number of stage-specific enhancers with evidence of positive selection in each development stage is indicated inside each bar. Only enhancers with at least 2 substitutions were used for this analysis. A. *The p-values from pairwise Fisher’s exact tests between TP3 and TP1, TP2, TP4, TP5 are 1.08 × 10^−15^, 1.01 × 10^−33^, 6.75 × 10^−25^, 9.94 × 10^−9^ respectively.* B. The p-values from pairwise Fisher’s exact tests between TP3 and TP1, TP2, TP4, TP5 are 5.04 × 10^−25^, 1.47 × 10^−46^, 4.57 × 10^−46^, 1.64 × 10^−7^ respectively.

To check whether our test could be biased by mutation patterns, we analyzed the rates of all possible substitutions and their corresponding deltaSVM (Liu and Robinson-Rechavi 2020), in each development stage. We split all substitutions into two categories: substitutions on CpG, and substitutions not on CpG. Overall, as expected, we found that the transition rate is much higher than the transversion rate, but we didn’t find any trend for specific substitution types to strengthen or weaken deltaSVM (Supplemental Fig. S4-8). We also checked whether neighboring substitutions (dinucleotide substitutions) have a general tendency to change deltaSVM in the same direction. Indeed, this is the case (Supplemental Fig. S9), suggesting that our test could be liberal or conservative for dinucleotide substitutions, depending on the direction of deltaSVM. Finally, to check whether dinucleotide substitutions and mutation bias affect the pattern we found, we excluded dinucleotide substitution sequences from all binding site, and we integrated the transition and transversion rate (3:1, estimated from Supplemental Fig. S4-8) into our null model. With these controls, we found a very consistent pattern to the original analysis (Supplemental Fig. S10). Thus our results are robust to the main known mutational biases.

It has been suggested that evolutionary pressure on enhancers varies in a tissue-specific manner across development (Nord et al. 2013). To check whether lower positive selection in the phylotypic period holds true at a tissue level, we assigned each enhancer from TP2 (6-8h) and TP3 (10-12h) into three categories (muscle, neuron and others) based on a recent lineage-resolved chromatin accessibility dataset in *D. melanogaster* (Reddington et al. 2020 and see Methods). This dataset contains muscle and neuron specific distal regulatory elements (putative enhancers) across four development stages, including two overlapping with TP2 and TP3 of our study. The pattern is consistent in each category (Supplemental Fig. S11), with lower positive selection at the phylotypic period and no strong difference between neural and muscle lineages. Thus the result is not driven by a specific pattern in one tissue, at least at this resolution.

## Discussion

Based on DNase I hypersensitive sites (DHS) in two distant *Drosophila* species across multiple matched embryonic stages, we identified a set of highly conserved stage-specific developmental enhancers. There is a higher proportion of conserved enhancers at the phylotypic period than at other embryonic stages, suggesting that conserved expression in the phylotypic period can be at least partly explained by conservation in gene regulation (Fig. 6). This provides a regulatory basis for the hourglass expression pattern. It was suggested that pleiotropic constraints might play important roles in the conservation of phylotypic stage (Duboule 1994; Raff 1996; Hu et al. 2017; Liu and Robinson-Rechavi 2018a). Since we found the highest conservation at the phylotypic stage for both stage specific enhancers and all enhancers (Fig. 2C and Supplemental Fig. S3), it’s less likely that the evolution of enhancers in the phylotypic stage was constrained by ontogenetic pleiotropic effects (e.g., enhancers used in multiple stages). However, we cannot rule out that our estimate of stage specificity might lack power, as it is based on a sampling of five embryonic stages. If data were available for more stages, with a finer temporal resolution, we might find that TP3 (phylotypic stage) specific enhancers are also used in neighboring developmental stages. In addition, the pleiotropic effects might be manifested at an anatomical level. For example, TP3 specific enhancers might be active in more cell types or tissues, and hence have higher pleiotropy not measured here. While enhancer conservation can be due to stronger purifying selection, it can also result from weaker positive selection. Here, for the first time, we provide evidence that the higher enhancer conservation at the phylotypic period can be explained in part by the latter (Fig. 6). This is consistent with similar results at the protein sequence level (Liu and Robinson-Rechavi 2018b; Coronado-Zamora et al. 2019). While both gene regulation and protein might be constrained at mid-development by lower embryo modularity, the new pattern we find for enhancer evolution can be more directly linked to previous observations of conservation of gene expression and of embryo morphology. Overall, we found that regulatory elements are more conserved in the phylotypic period, and less accessible to adaptive evolution. Thus, the phylotypic period can be regarded as an evolutionary regulatory lockdown.

**Figure 6:**
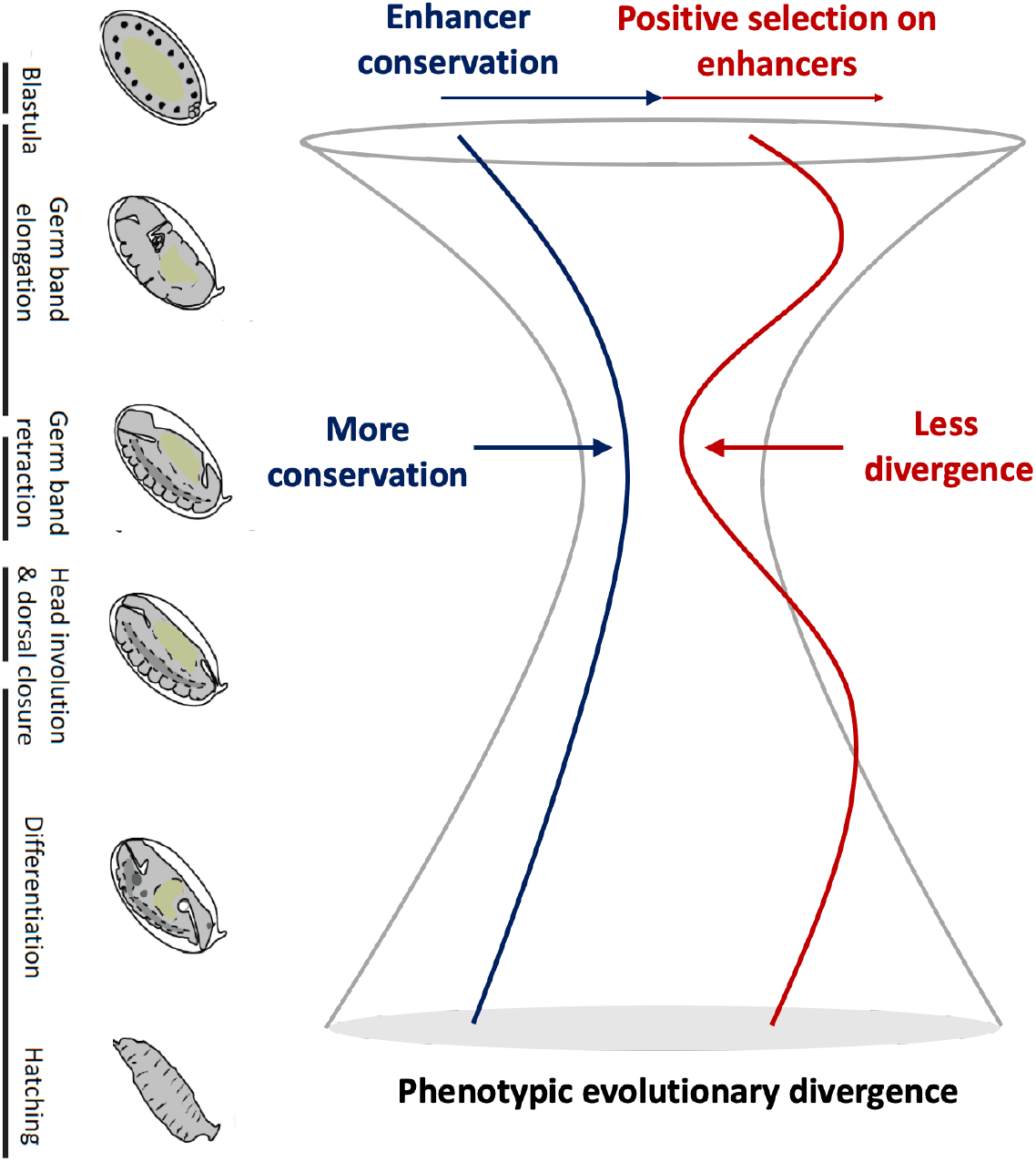
A simple model of the evolutionary forces on gene regulation for the hourglass pattern. Embryo images of D. melanogaster adapted from Levin et al. (2016).

Our results indicate that widespread positive selection shaped the evolution of developmental enhancers, especially of early and late embryonic enhancers. This signature does not appear to be driven by a specific tissue, at least at the level of the two major lineages examined (muscle and neuron). Higher adaptation in late embryonic stages could be due to the greater diversity of challenges to which natural selection needs to respond in the next stages of larva and juvenile development, compared to early and mid-embryogenesis. This fits with Darwin’s “selection opportunity” (Liu and Robinson-Rechavi 2018b). Adaptation in the early embryo is less expected. Whereas early developmental proteins are influenced by the maternal contribution, which could be impacted by selection on reproduction, early embryonic enhancers are specific to the zygote. While there is evidence that the evolution of *cis*-regulatory elements is the main driver of morphologic diversity (Gompel et al. 2005; McGregor et al. 2007; Wray 2007; Jeong et al. 2008), for most changes in early embryogenesis there are no clear consequences on adult morphology (Kalinka and Tomancak 2012). This raises the question of why there are so many adaptive enhancer changes in early embryogenesis. One explanation, proposed by Kalinka and Tomancak (2012), is that much of the variation in early embryogenesis results from adaptation to diverse ecological circumstances. For example the evolution of long germ segmentation in some insects (Liu and Kaufman 2005) might play a role in shortening the embryonic development time, which is likely an adaptive strategy to a particular ecological niches (Kalinka and Tomancak 2012). Another explanation is that adaptive evolution of maternally contributed *trans* factors might drive rapid compensatory evolution of early zygotic enhancers. It should also be noted that the maxima of positive selection are at the second and fourth time points, i.e. not at the first and last. An interesting speculation is that the stages with the most positive selection on enhancers are those where the most cell differentiation is occurring, but this is highly speculative in the present state of our data and our knowledge, and while it fits to our later time-points (TP4 and TP5), it does not to the early ones (TP1, TP2).

While we show a role for lower adaptive evolution in the regulation of more conserved expression at the phylotypic period, several recent studies also found evidence of developmental constraints, both purifying selection and mutational robustness. For example, in order to mostly eliminate the influence of positive selection on gene expression evolution, Zalts and Yanai (2017) quantified expression variability of twenty *C. elegans* mutation accumulation strains throughout embryogenesis. They found that the nematode phylotypic period has lower expression variability, indicating that purifying selection can contribute to the hourglass model of expression evolution. These results are also compatible with a role of mutational robustness. In *D. melanogaster*, we have recently compared the expression variability of isogenic single embryo transcriptomes across development and found lower variability at the phylotypic period, suggesting that genes expressed at this stage are intrinsically less sensitive to perturbations on gene expression (Liu et al. 2020). Here, we found a much higher number of enhancers at the phylotypic period, which suggests more redundancy and thus higher regulatory robustness in gene expression. This could also contribute to mutational robustness (Cannavò et al. 2016) and thus lower expression divergence. Overall, the low expression divergence at the phylotypic period seems to have been shaped by the interplay of purifying selection, positive selection, and mutational robustness.

## Methods

### DNase-seq on *D. melanogaster* and *D. virilis* embryos

The first three time-points were from our published study (Peng et al. 2019; with EMBL-EBI ArrayExpress accession number E-MTAB-3797). Here we supplemented those by two additional time-points in each species to extend our time-course beyond the phylotypic period (Fig. 1A). Stage matched *D. melanogaster* and *D. virilis* embryos (stages defined by Khoueiry et al. (2017)) were collected and DNase-seq performed as described previously (Khoueiry et al. 2017). Two biological replicates were generated for each species at every stage.

### DNase-seq sample processing

All analysis were performed in the Galaxy platform (Afgan et al. 2018). Raw paired-end reads were first trimmed with Trim Galore (https://github.com/FelixKrueger/TrimGalore, Galaxy Tool Version 0.4.3.1) and reads were clipped to maximum of 94 bases using Trimmomatic (Bolger et al. 2014) (Galaxy Tool Version 0.36.6). Then the trimmed reads were mapped to the *D. melanogaster* genome (Flybase Assembly 6 version: dm6) and to the *D. virilis* genome (Flybase-R1.2 assembly version: droVir3) respectively by using Bowtie2 (Langmead and Salzberg 2012) with standard parameters (Galaxy Tool Version 2.3.4.2). Multi-mapping reads were discarded and duplicates were removed using MarkDuplicates (Galaxy Tool Version 2.7.1.1) from the Picard suite https://github.com/broadinstitute/picard. For peak calling, we used MACS2 (Zhang et al. 2008) with standard parameters (MACS2 Galaxy Tool Version 2.1.1.20160309.5). We derived peaks using 5% Irreproducible Discovery Rate (IDR, Galaxy Tool Version 2.0.3) threshold for biological replicates.

### Genome coordinates translation

Since *D. melanogaster* and *D. virilis* are highly divergent, the *D. virilis (D. melanogaster)* peak coordinates were translated to *D. melanogaster (D. virilis)* genome coordinates by using the pslMap (Zhu et al. 2007), as suggested by Khoueiry et al. (2017).

### Sequence alignment files

The pairwise whole genome alignments between *D. melanogaster* and *D. simulans* or *D. yakuba* were downloaded from Haeussler et al. (2019) http://hgdownload.soe.ucsc.edu/downloads.html (accessed in December, 2018).

### Single Nucleotide Polymorphism (SNP) data

Over 4.8 million SNPs for 205 *D. melanogaster* inbred lines were downloaded from the Drosophila Genetic Reference Panel (DGRP) http://dgrp2.gnets.ncsu.edu/ (Huang et al. 2014, accessed in December, 2018).

### In silico mutagenesis for detecting positive selection on enhancers

1. Training the gapped k-mer support vector machine (gkm-SVM) gkm-SVM is a method for predicting regulatory DNA sequence by using *k*-mer frequencies (Ghandi et al. 2014). For the gkm-SVM training, we followed the same approaches as Lee et al. (2015). Firstly, we defined a positive training set and its corresponding negative training set. The positive training set is stage-specific enhancers. The negative training set is an equal number of sequences randomly sampled from the genome with matched length, GC content and repeat fraction as in the positive training set. This negative training set was generated by using “genNullSeqs”, a function of the gkm-SVM R package (Ghandi et al. 2016). Then, we trained a gkm-SVM with default parameters except −l=10 (meaning we use 10-mers as feature to distinguish positive and negative training sets). The classification performance of the trained gkm-SVM was measured by using receiver operating characteristic (ROC) curves with fivefold cross-validation. The gkm-SVM training and cross-validation were achieved by using the “gkmtrain” function of “LS-GKM” (Lee 2016) (also see https://github.com/Dongwon-Lee/lsgkm).
2. Testing the stage specificity of gkm-SVM To test the performance of gkm-SVM trained in stages other than the focal stage (e.g., trained TP2, TP3, TP4, TP5 to predict in TP1), we first scored both positive and negative training sets in the focal stage by using the gkm-SVM from other stages. We used the “gkmpredict” function of “LS-GKM”. Then, the ROC curve was used to evaluate the prediction performance.
3. Generating SVM weights of all possible 10-mers The SVM weights of all possible 10-mers were generated with the “gkmpredict” function of “LS-GKM”. A positive value means increasing chromatin accessibility, a negative value means decreasing accessibility, and value close to 0 means no impact on chromatin accessibility (the function measured in this case).
4. Infering ancestor sequence The ancestor sequence was inferred from sequence alignment between *D. melanogaster* and *D. simulans* by using *D. yakuba* as an outgroup.
5. Calculating deltaSVM We calculated the sum of weights of all 10-mers for ancestor sequence and focal sequence respectively. The deltaSVM is the sum of weights of the focal sequence minus the sum of weights of the ancestor sequence. A positive deltaSVM indicates substitutions increasing the chromatin accessibility in the focal sequence, and vice versa.
6. Generating Empirical Null Distribution of deltaSVM Firstly, we counted the number of substitutions between each ancestor sequence and focal sequence. Then, we generated a random pseudo-focal sequence by randomly introducing the same number of substitutions to the ancestor sequence. Finally, we calculated the deltaSVM between the pseudo-focal sequence and the ancestor sequence. We repeated this process 10000 times to get 10000 expected deltaSVMs.
7. Calculating *p-*value of deltaSVM The *p-*value was calculated as the probability that the expected deltaSVM is higher than the observed deltaSVM. The *p-*value can be interpreted as the probability that the observed deltaSVM could arise by chance under the assumptions of the randomisation.

### Estimate substitution rate

The substitution rate, for example C -> T, was estimated as the number of C -> T divided by the number of nucleotide C in the ancestor sequence.

### Definition of conserved and non-conserved enhancers

We split stage-specific enhancers into two categories: conserved and non-conserved. A *D. melanogaster* TP*i* specific enhancer whose orthologous region in *D. virilis* overlaps at least 1bp with a *D. virilis* enhancer is defined as conserved. All other *D. melanogaster* TP*i* enhancers are defined as non-conserved.

### Muscle and neuron specific enhancer assignment

We first downloaded muscle and neuron specific enhancers (DNase-seq identified peaks greater than 500bp from an annotated transcriptional state site (TSS)) from Reddington et al. (2020). This study included DNase-seq in isolated muscle and neural lineages across four stages (4-6h, 6-8h, 8-10h, 10-12h) of *Drosophila melanogaster* embryogenesis. Then, based on these tissue specific enhancers, we split our whole embryo identified TP2 (6-8h) specific and TP3 (10-12h) specific enhancers into three categories: muscle specific enhancers, neuron specific enhancers, and remaining enhancers. A TP2 specific enhancer which overlaps at least 1bp with a muscle (resp. neuron) specific enhancer is defined as a TP2 muscle (resp. neuron) specific enhancer. All other enhancers are defined as “remaining” enhancers. The same was applied to TP3 specific enhancers.

## Data access

DNase-seq data for the last two time points (TP4 and TP5) in each species was deposited to EMBL-EBI ArrayExpress with accession number E-MTAB-9480. Data files and analysis scripts are available on GitHub: (https://github.com/ljljolinq1010/Chromatin-accessibility-evolution-during-Drosophila-embryogenesis)

## Acknowledgements

We thank the EMBL Genomics Core Facilities for support, and David Garfield and Gunter Wagner for critical reading of an earlier version of the manuscript. We thank members of the Furlong and Robinson-Rechavi labs for helpful discussions. We thank Irepan Salvador-Martínez, Axel Visel and Matt Benton for comments on a previous version in preprint. Part of the computations were performed at the Vital-IT (http://www.vital-it.ch) Center for high-performance computing of the SIB Swiss Institute of Bioinformatics. EEF is supported by the Marie Curie EvoNet ITN and the DFG grant FU 750; JL and MRR are supported by Swiss National Science Foundation grant 31003A_173048.

Author contributions: EEF initiated the project, and supervised all experiments. RV, PK and JR performed all experiments. CG performed raw data analysis and DNase-seq peak calling. JL designed the detailed study with input from MRR. JL performed all downstream bioinformatics analyses. JL and MRR interpreted the results with input from all other authors. JL wrote the first draft of the paper. JL, EEF and MRR finalized the paper with input from all authors.

