## supplementary figures for "The hourglass model of evolutionary conservation during embryogenesis extends to developmental enhancers with signatures of positive selection"

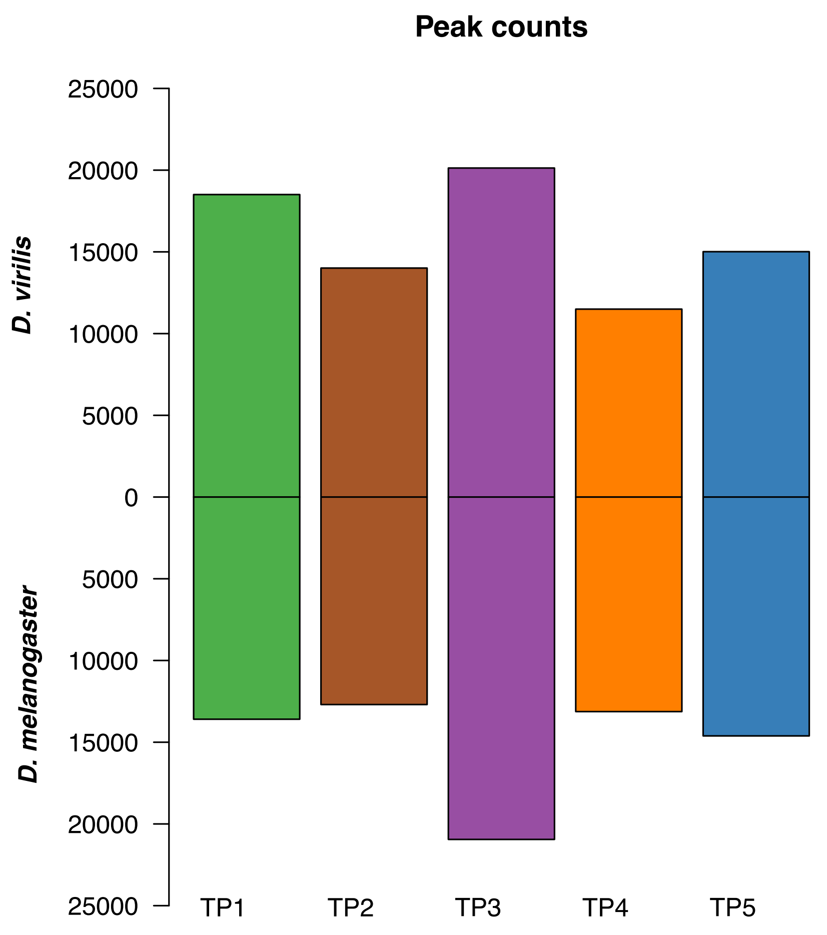


**Figure S1: The number of significant peaks across development**

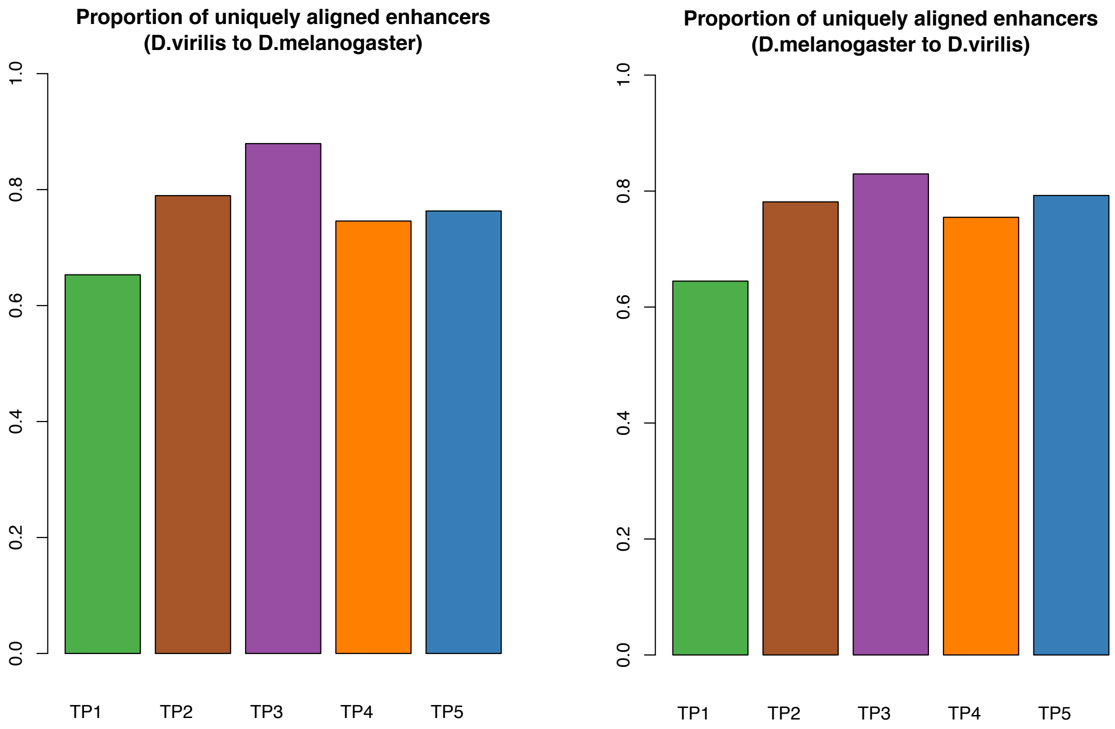


**Figure S2: The proportion of peaks identified in one species that with a one-to-one orthologous region in the other species.**

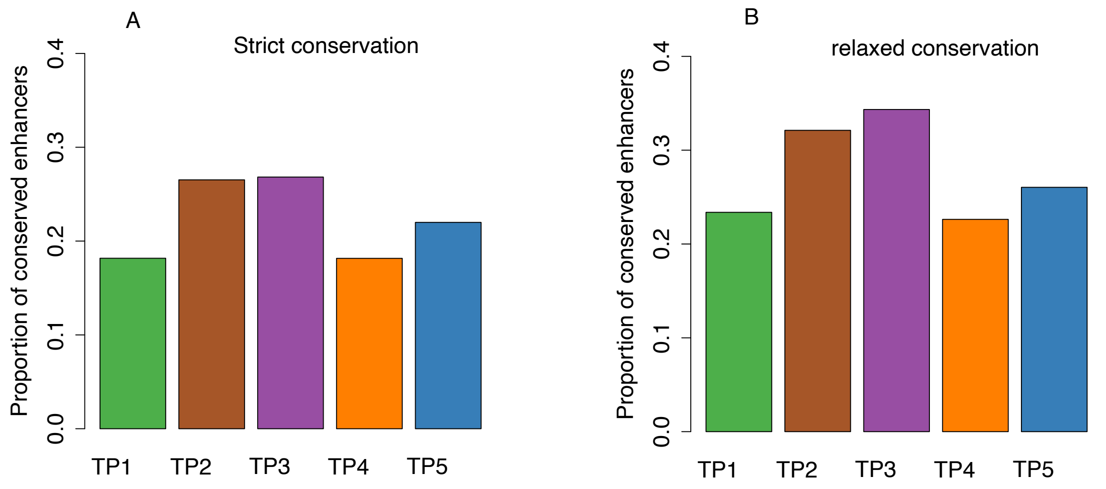


**Figure S3 : Proportion of conserved enhancers at each development stage.**

For each stage, we used all enhancers identified in this stage, not only the stage specific ones.

A. Here the conservation means there is at least 1bp overlap between enhancers in the two species. The p-values from pairwise Fisher's exact tests between TP3 and TP1, TP2, TP4, and TP5 are 7.86e-42, 0.71, 2.57e-31, 2.03e-11 respectively.

B. Here the conservation means the distance between enhancers in the two species must be smaller than 1kb, not necessarily overlap. The p-values from pairwise Fisher's exact tests between TP3 and TP1, TP2, TP4, TP5 are 3.63e-47, 0.0138, 1.04e-40, 4.21e-23 respectively.

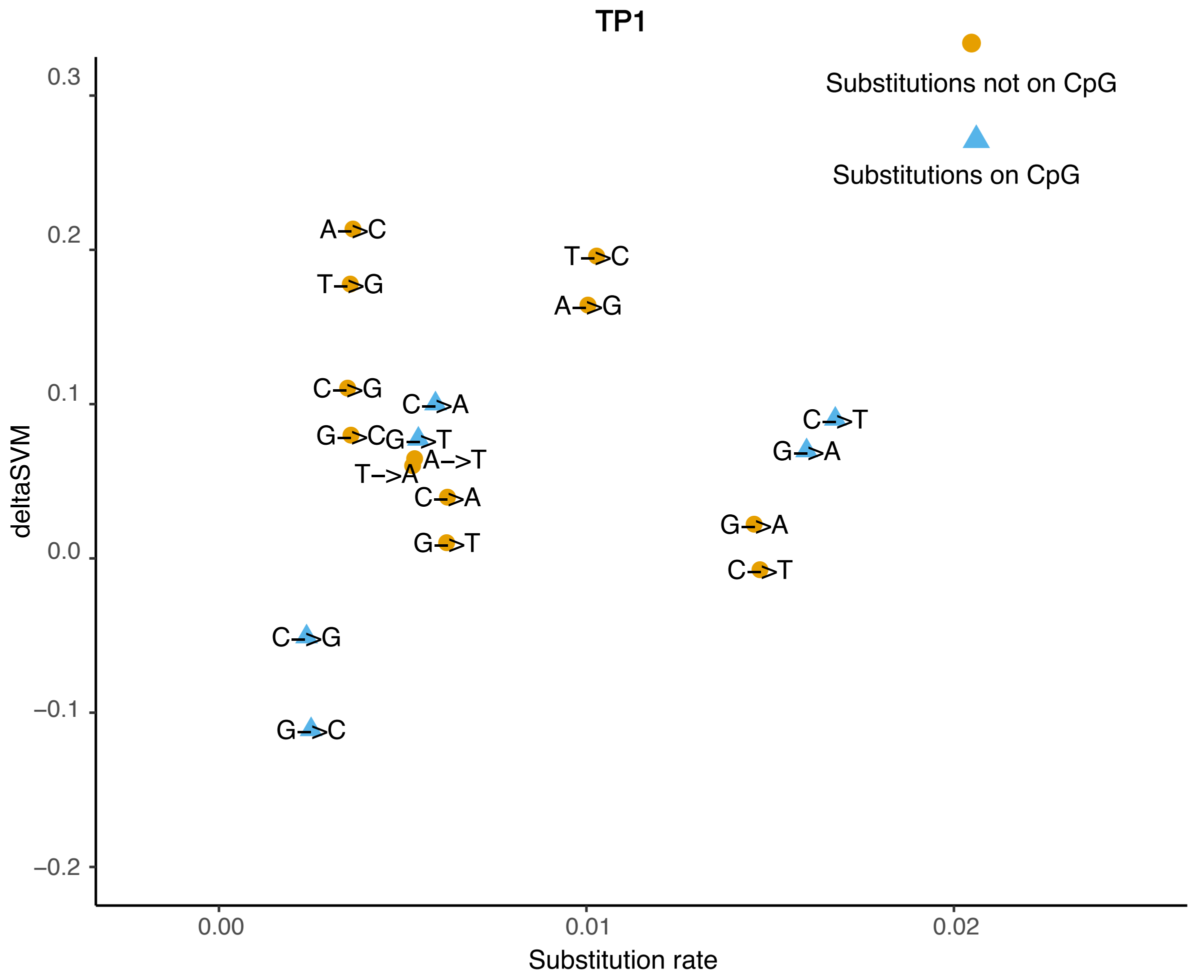


**Figure S4: Substitution type, substitution rate and deltaSVM relationship for the first stage.**

The x-axis is the substitution rate of different types of substitutions, the y-axis is the median deltaSVM of different types of substitutions.

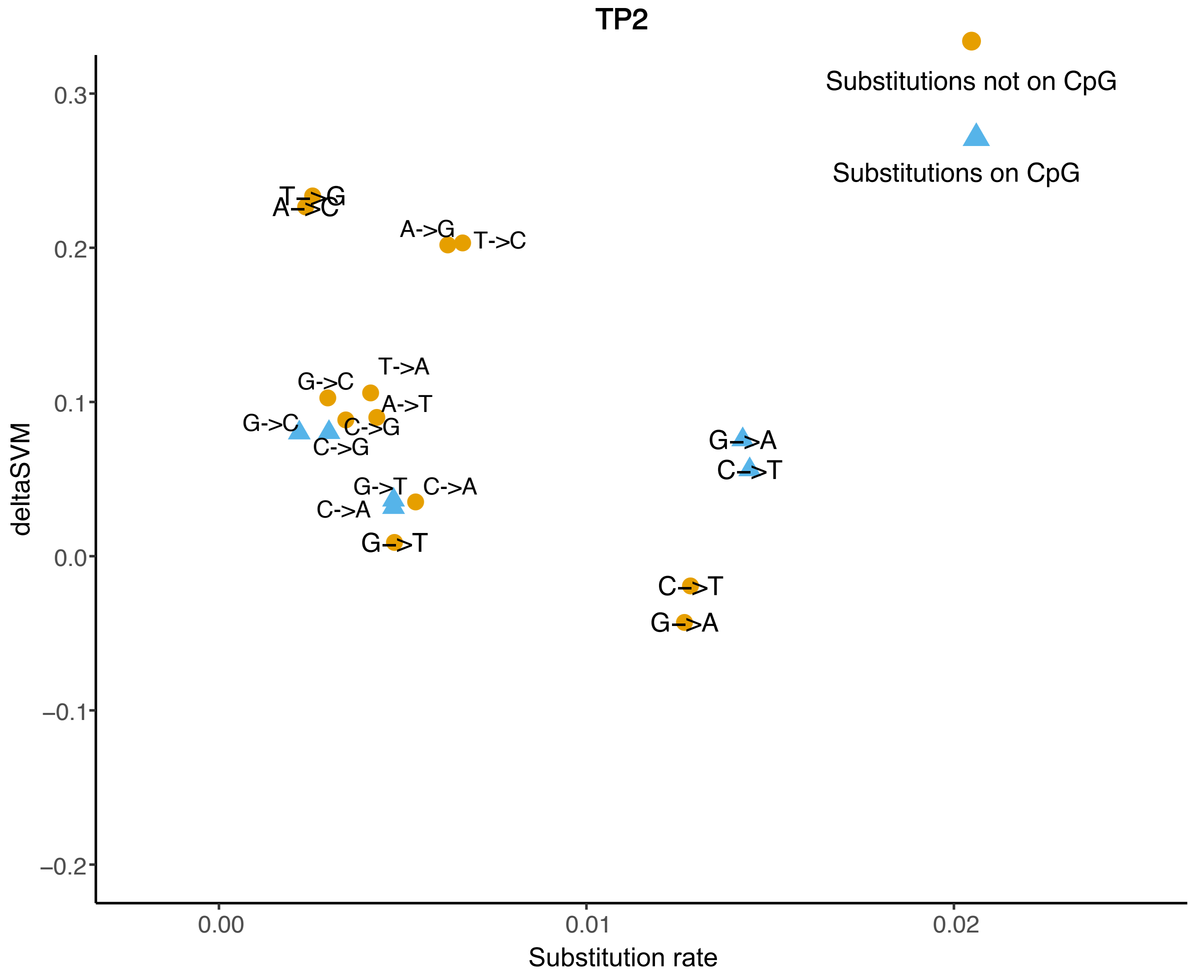


**Figure S5: Substitution type, substitution rate and deltaSVM relationship for the second stage.**

The x-axis is the substitution rate of different types of substitutions, the y-axis is the median deltaSVM of different types of substitutions.

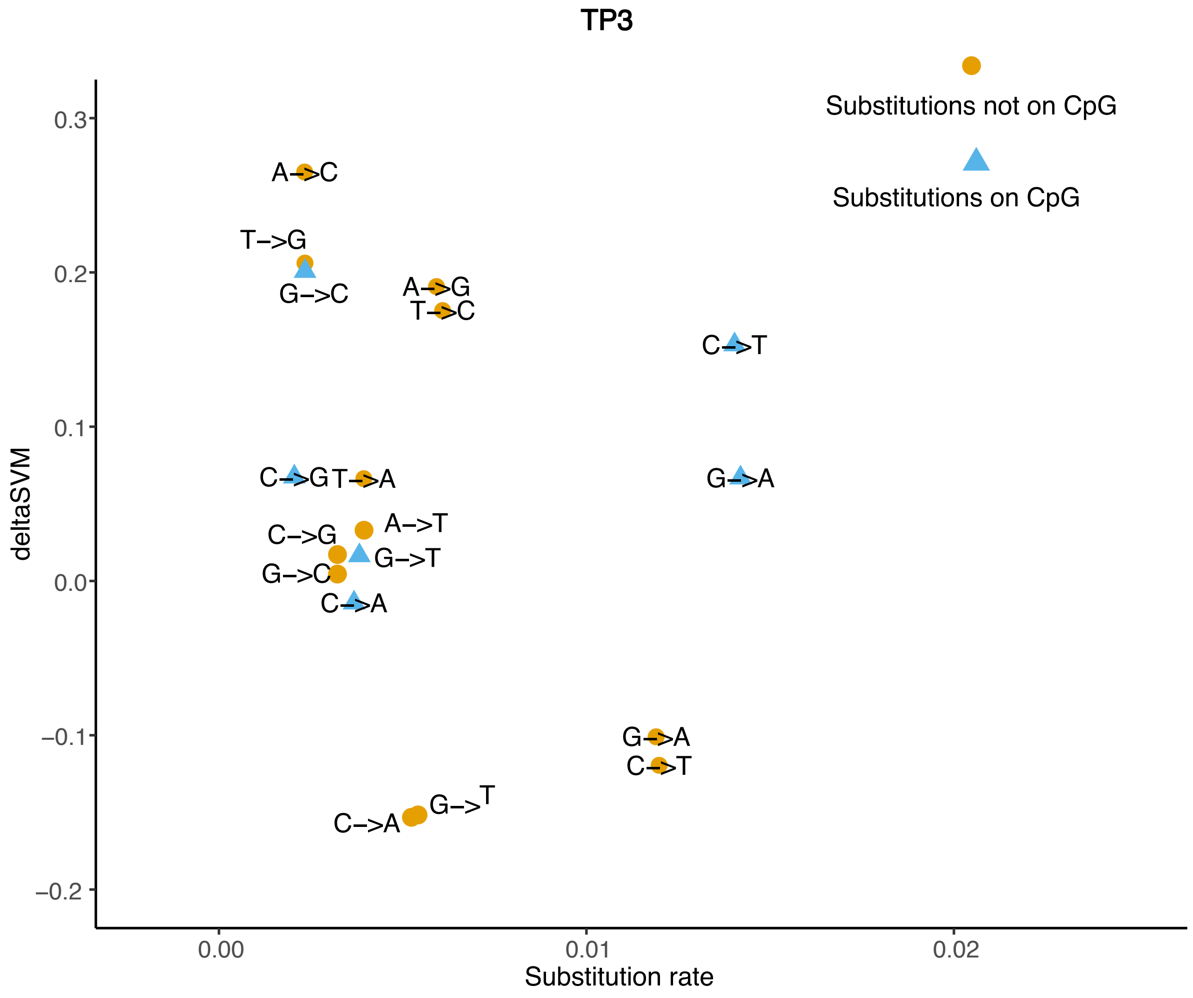


**Figure S6: Substitution type, substitution rate and deltaSVM relationship for the third stage.**

The x-axis is the substitution rate of different types of substitutions, the y-axis is the median deltaSVM of different types of substitutions.

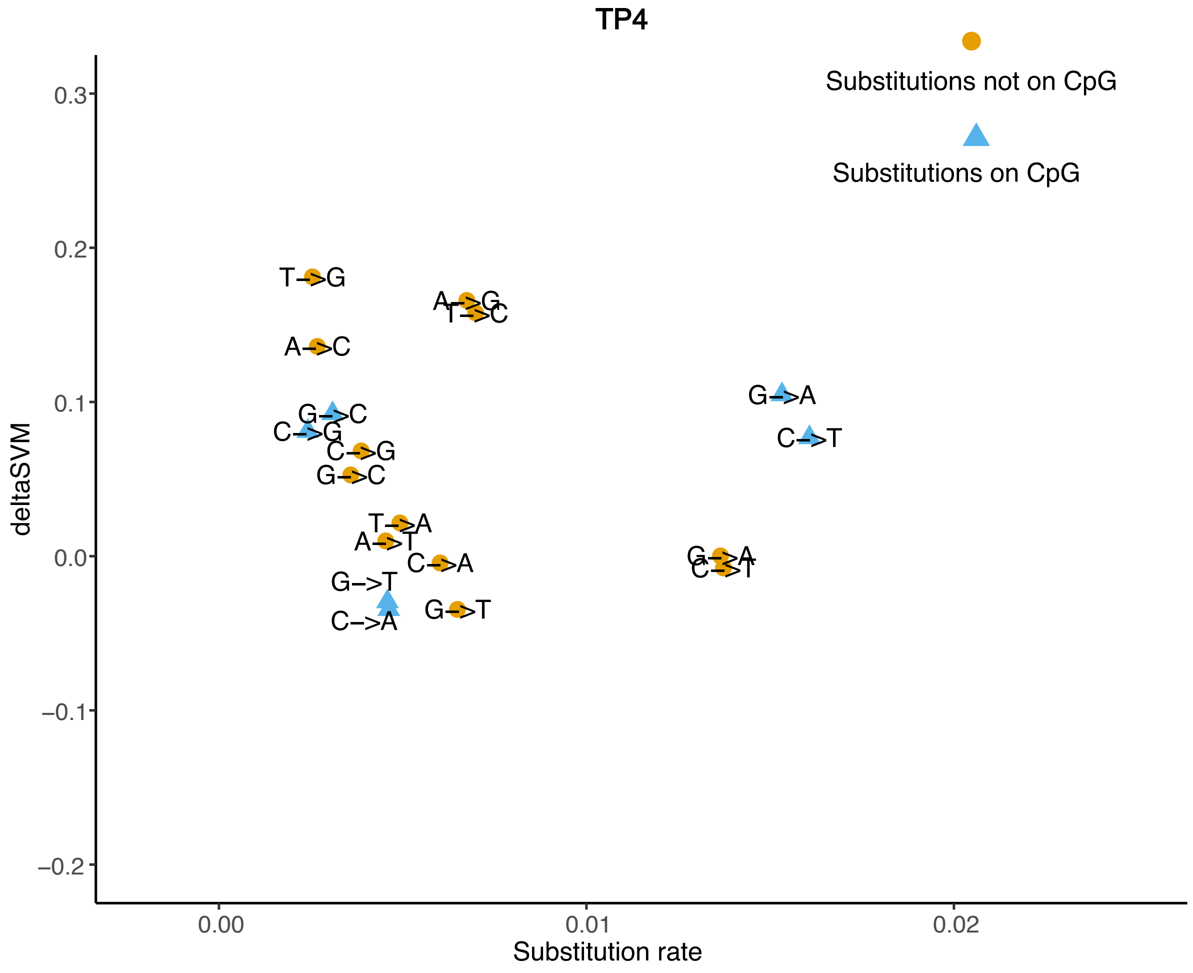


**Figure S7: Substitution type, substitution rate and deltaSVM relationship for the fourth stage.**

The x-axis is the substitution rate of different types of substitutions, the y-axis is the median deltaSVM of different types of substitutions.

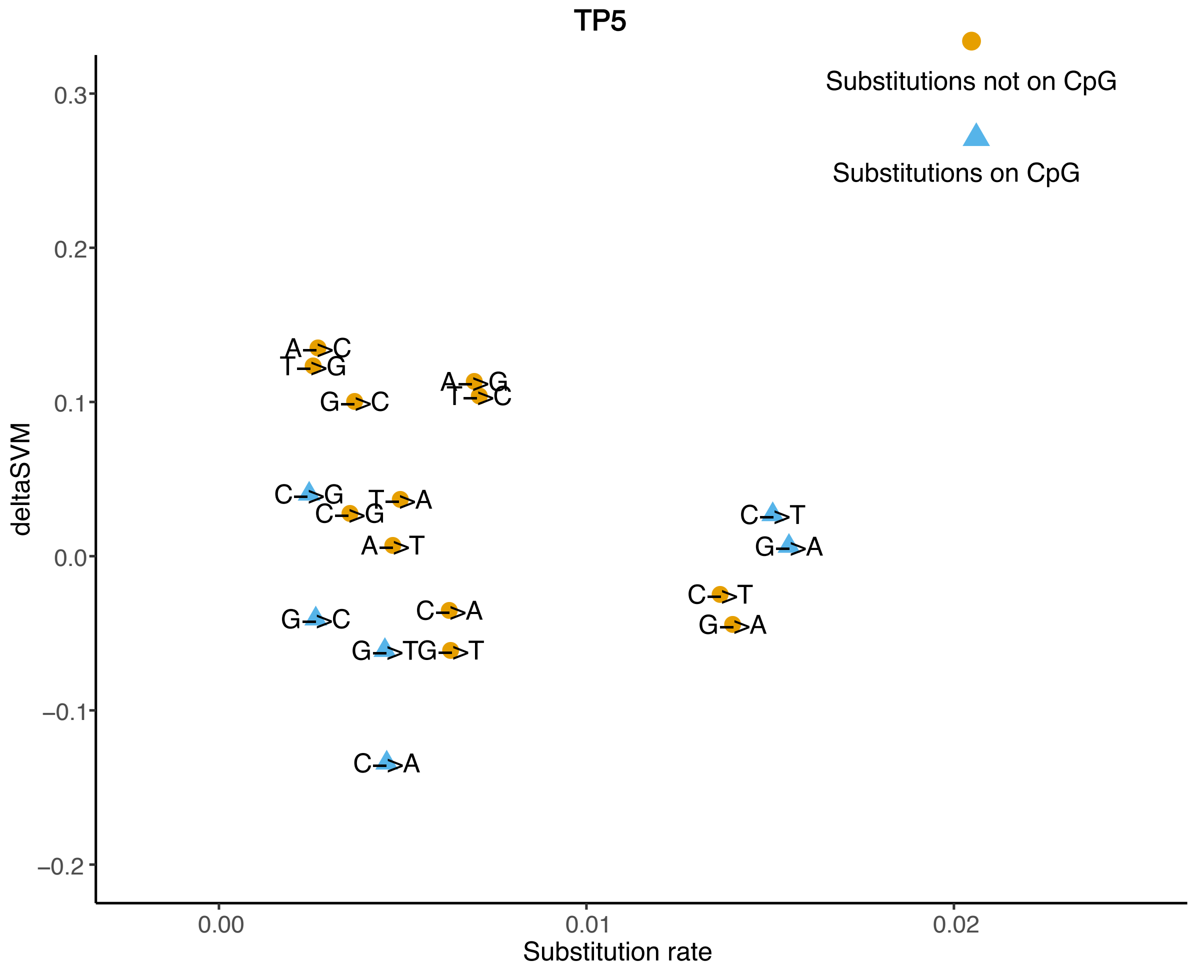


**Figure S8: Substitution type, substitution rate and deltaSVM relationship for the fifth stage.**

The x-axis is the substitution rate of different types of substitutions, the y-axis is the median deltaSVM of different types of substitutions.

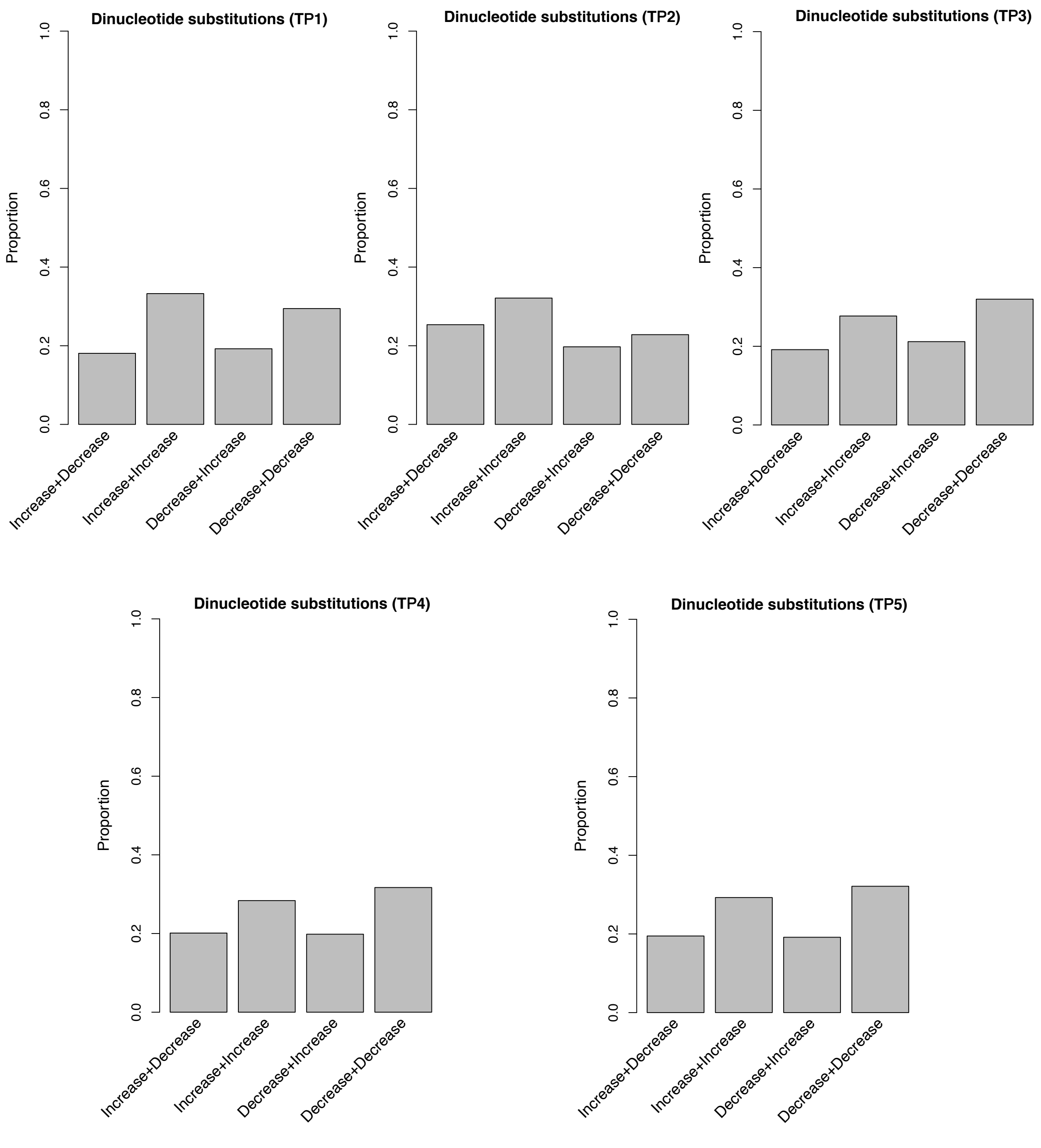


**Figure S9: Direction of binding affinity change of dinucleotide substitutions**

The label below the bottom of each bar indicates the direction of binding affinity change of two neighboring substitutions. For example, Increase+Decrease indicates the first substitution increases the binding affinity, but the second substitution decreases the binding affinity.

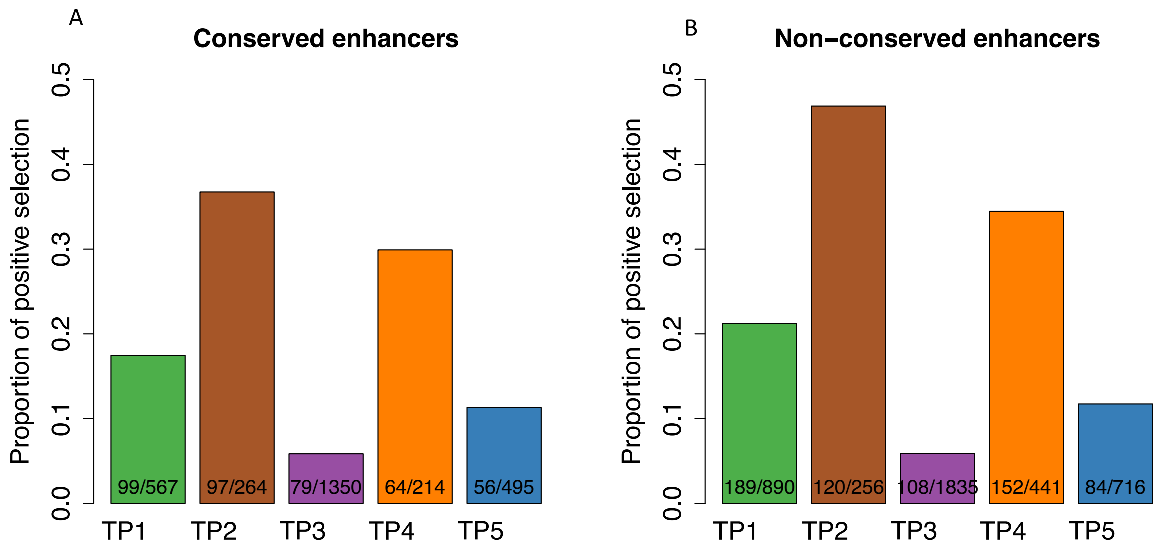


**Figure S10: The proportion of enhancers with evidence of positive selection from the new analysis.**

In the new analysis, we excluded all CpG sequences and dinucleotide substitution sequences, and controlled the transition and transversion rate. Positive sites are enhancers with evidence of positive selection (deltaSVM qvalue < 0.05). The number of stage specific enhancers and the number of stage specific enhancers with evidence of positive selection in each development stage is indicated inside each bar.

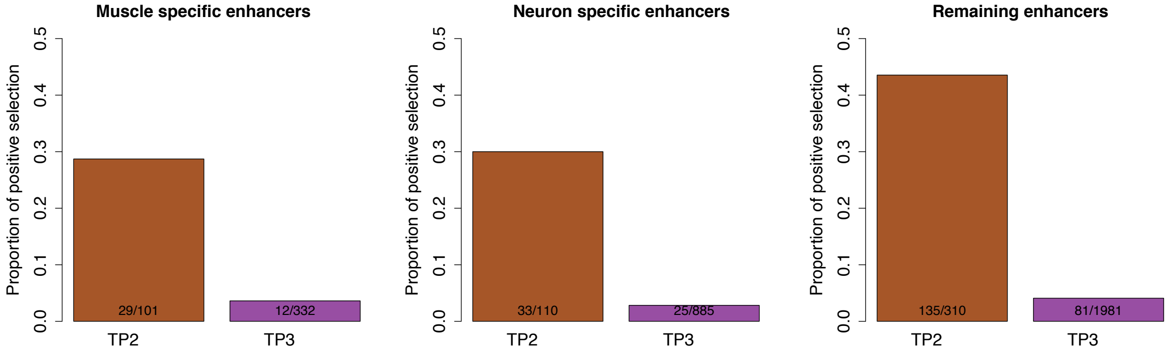


**Figure S11: The proportion of enhancers with evidence of positive selection for tissue specific enhancers.**

Positive sites are enhancers with evidence of positive selection (deltaSVM qvalue $< 0.05$). The number of stage specific enhancers and the number of stage specific enhancers with evidence of positive selection in each development stage is indicated inside each bar.
